# Genome diversity of *Leishmania aethiopica*

**DOI:** 10.1101/2023.01.17.524362

**Authors:** Amber Hadermann, Senne Heeren, Ilse Maes, Jean-Claude Dujardin, Malgorzata Anna Domagalska, Frederik Van den Broeck

## Abstract

*Leishmania aethiopica* is a zoonotic Old World parasite transmitted by Phlebotomine sand flies and causing cutaneous leishmaniasis in Ethiopia and Kenya. Despite a range of clinical manifestations and a high prevalence of treatment failure, *L. aethiopica* is the most neglected species of the *Leishmania* genus in terms of scientific attention. Here, we explored the genome diversity of *L. aethiopica* by analyzing the genomes of twenty isolates from Ethiopia. Phylogenomic analyses identified two strains as interspecific hybrids involving *L. aethiopica* as one parent and *L. donovani* and *L. tropica* respectively as the other parent. High levels of genome-wide heterozygosity suggest that these two hybrids are equivalent to F1 progeny that propagated mitotically since the initial hybridization event. Analyses of allelic read depths further revealed that the *L. aethiopica* - *L. tropica* hybrid was diploid and the *L. aethiopica* - *L. donovani* hybrid was triploid, as has been described for other interspecific *Leishmania* hybrids. When focusing on *L. aethiopica*, we show that this species is genetically highly diverse and consists of both asexually evolving strains and groups of recombining parasites. A remarkable observation is that some *L. aethiopica* strains showed an extensive loss of heterozygosity across large regions of the chromosomal genome, which likely arose from gene conversion/mitotic recombination. Hence, our prospection of *L. aethiopica* genomics revealed new insights into the genomic consequences of both meiotic and mitotic recombination in *Leishmania*.

## INTRODUCTION

*Leishmania* is a vector-borne parasite causing leishmaniasis in humans and animals. Depending on the *Leishmania* species, the disease can present itself in various clinical representations, ranging from cutaneous to visceral leishmaniasis. *Leishmania aethiopica* is a zoonotic Old World species transmitted by sand flies of the *Phlebotomus* genus, and is known to cause local (LCL), diffuse (DCL) and the occasional mucocutaneous leishmaniasis (MCL). Where LCL will present as self-healing lesions at the place of inoculation, DCL appears as non-self-healing lesions widespread over the whole body and MCL at the mucosal membranes (i.e. nose, mouth and throat) [1–4]. This species is endemic in Ethiopia and the highlands of Kenya, with respectively 1,402 and 398 reported cases of cutaneous leishmaniasis in 2020 [3]. However, the total burden of *L. aethiopica-associated* disease is difficult to estimate since most of the infections will not result in disease, and if they do, they often remain unreported due to the risk of stigmatization [2]. In addition to being a species that clinically presents three forms of CL, *L. aethiopica* has a high prevalence of treatment failure that remains poorly understood [3]. Finally, *L. aethiopica* field isolates bear the endosymbiotic and immunogenic double-stranded *Leishmania* RNA virus, which may have potential implications on disease severity [5]. Despite these observations of biomedical and epidemiological relevance, research on *L. aethiopica* is almost non-existent, leaving large gaps in the knowledge on the biology of this most neglected *Leishmania* species.

Genome diversity studies allow understanding the population dynamics and biology of *Leishmania* parasites, revealing insights into e.g. the epidemic history of the deadly *L. donovani* in the Indian subcontinent [6], the colonization history of *L. infantum* in the Americas [7], the Pleistocene origin of the Andean *Leishmania* parasites [8] and the genetic consequences of hybridization [9]. Unfortunately, studies on the genetic diversity of *L. aethiopica* are scanty and limited to analyses of microsatellite genotyping, isoenzyme analysis, fragment length polymorphism analyses and/or single gene sequencing [10–13]. These molecular studies demonstrated that the species *L. aethiopica* is genetically very heterogeneous despite its restricted geographic distribution [10–12] and suggested the existence of a *L. aethiopica/L. donovani* hybrid (MHOM/ET/94/ABAUY) [13]. Such findings indicate that *L. aethiopica*-related disease may be caused by a genetically diverse and recombining species, which may have consequences towards the clinical management and epidemiology of CL in East Africa.

Here, we present the first genome diversity study of *L. aethiopica* to gain a better understanding of the evolutionary history and population biology of this species. We generated genomes for a total of 28 *Leishmania* isolates, including 20 *L. aethiopica* isolates from Ethiopia, and complemented our dataset with publicly available genomes from *L. donovani, L. infantum*, *L. tropica* and *L. major* for comparative purposes. Population genomic and phylogenomic analyses provide genomic confirmation of interspecific hybridization including *L. aethiopica* as one of the two parental species, and confirm that *L. aethiopica* is genetically highly diverse. Our prospection of *L. aethiopica* genomics will promote future studies on the genomic basis of treatment failure and clinical outcome.

## METHODS

### Dataset of genomic sequences

A total of 28 *Leishmania* isolates were sequenced within the context of this study (Supp. Table 1), including 18 *L. aethiopica* isolates collected in Ethiopia from 1959 to 1994 and previously typed by the Centre National de Référence des *Leishmania* (Montpellier, France) and the WHO International *Leishmania* Reference Center (London School of Hygiene and Tropical Medicine, London, United Kingdom). For two isolates (GERE and KASSAYE), a derived clone was sequenced (Supp. Table 1). Our dataset also included isolate MHOM/ET/72/L100 and its derived clone L100cl1; the L100 strain is the WHO reference strain that represents the same strain as the one used to create the reference genome (https://tritrypdb.org/tritrypdb/app/record/dataset/NCBITAXON_1206056#description). In addition, we included isolate MHOM/ET/94/ABAUY (here-after referred to as LEM3469) and its five derived clones (Supp. Table 1), in order to confirm hybridization between *L. aethiopica* and *L. donovani* and investigate its genomic consequences. All isolates were sequenced (150bp paired-end) on the Illumina sequencing platform of GenomeScan, Leiden, The Netherlands. For comparative purposes, we included publicly available sequences from four Old World *Leishmania* species (Supp. Table 1): *L. donovani* (n=5)*, L. infantum* (n=1), *L. major* (n=1) and *L. tropica* (n=1) [14–16]. Altogether, this provided us with a total of 36 genomes for downstream analyses.

### Bioinformatic analyses

All sequences were mapped against the *L. aethiopica’s* reference genome L147 (available on tritrypdb.org as TriTrypDB-54_LaethiopicaL14710) using BWA [17]. Resulting SAM-files were converted to BAM-files using SAMtools [18]. All duplicates were marked using the GATK (Genome Analysis ToolKit) software [19]. BAM-files were checked for mapping quality by examining flagstat files and coverage was calculated with SAMtools depth to determine the average mapped read depth across the whole genome. SNPs and INDELs were called twice with GATK HaplotypeCaller: once including all 36 *Leishmania* genomes and once including the 20 *L. aethiopica* isolates.

Single Nucleotide Polymorphisms (SNPs) and small insertions/deletions (INDELs) were separated with the GATK SelectVariants command. SNPs were hard filtered based on the GATK filter recommendations by G. Van der Auwera, including a mapping quality (MQ) larger than 40 and a quality by depth (QD) larger than 2, in combination with a filter that excludes SNPs within SNP clusters (defined by 3 SNPs in windows of 10bp) [20]. In addition, SNPs were retained only when the allelic depth per genotype (i.e. Format Depth, FMTDP) was larger than 5 and the genotype quality (GQ) was larger than 20 (when genotyping was done across all 36 *Leishmania* genomes) or larger than 40 (when genotyping was done on the 20 *L. aethiopica* genomes). SNPs were counted per isolate for the whole genome and per chromosome in Rstudio [21]. Alternate allele read depth frequencies were counted using the vcf2freq.py script (available at github.com/FreBio/mytools/blob/master/vcf2freq.py).

### Chromosomal and local copy number variation

Chromosomal and local copy number variation were studied for the 20 *L. aethiopica* isolates and the two interspecific hybrids L86 and LEM3469 based on the per-site read depth as obtained with SAMTools depth. To obtain haploid copy numbers (HCN) for each chromosome, the median chromosomal read depths were divided by the median genome-wide read depth. Somy variation was then obtained by multiplying HCN by two (assuming diploidy for all *L. aethiopica* isolates and L86; see results) or three (assuming trisomy for isolate LEM3469 and its derived clones; see results). To obtain local HCN, the median read depth in non-overlapping 2kb windows was divided by the median genome-wide read depth. These calculations were done using the depth2window.py script (available at github.com/FreBio/mytools/blob/master/depth2window.py).

Windows of reduced or increased HCN were identified by deducting the median HCN across 20 *L. aethiopica* genomes from the HCN as estimated per *Leishmania* genome. This results in a distribution of HCN centered around zero, allowing us to identify 2kb windows with increased (z-score > 5) or decreased (z-score < −5) HCN in each of the 20 *L.aethiopica* isolates and the two interspecific hybrids (LEM3469 and L86). Consecutive 2kb windows showing a significant increase/decrease in HCN were joined by averaging HCN across the consecutive windows, allowing us to detect larger copy number variants. Small deletions/amplifications (i.e. <= 6kb) that do not cover protein coding genes were ignored.

### Population genomic and phylogenomic analyses

The quality-filtered SNP VCF files were converted to the fasta format using the vcf2fasta.py script (available at github.com/FreBio/mytools/blob/master/vcf2fasta.py). To get an initial idea on the phylogenetic relationships between the isolates, a phylogenetic network was reconstructed with SplitsTree version 4.17.2 [22]. For the chromosomal networks, the chromosomal SNPs were selected using BCFtools view and converted into the fasta format for network analyses.

The population genomic structure of *L. aethiopica* was investigated after excluding putative near-identical isolates (see results) and after removing sites exhibiting high linkage disequilibrium (LD). Non-independent SNPs were removed in a pairwise manner (--indep-pairwise) with plink v1.9 [23] within 50 bp windows with a 10 bp step size for three different squared correlation coefficients (r^2^=0.3, 47,244 SNPs retained; r^2^=0.5, 85,725 SNPs retained; r^2^=0.7, 112,241 SNPs retained). ADMIXTURE v1.3 was run for *K* equals 1 to 5 along with a five-fold cross validation [23]. Principal component analysis (PCA) was done using the glPCA function within the Adegenet R-package [25]. Nucleotide diversity (π), Tajima’s D and Weir and Cockerham’s pairwise F_ST_ (mean and weighted), were calculated in non-overlapping windows of 50kb for the populations as inferred by ADMIXTURE using VCFtools v0.1.13 [24]. Loss-of-Heterozygosity (LOH) regions were identified by analyzing the distribution of heterozygous sites in non-overlapping 10kb windows as described elsewhere [25], using the following criteria: min number of SNPs = 1, max number of heterozygous sites allowed per 10kb window = 0, minimum number of contiguous 10kb windows = 4, maximum ⅓ of all 10 kb windows within a LOH region can be gap, max number of heterozygous sites allowed within gap = 2.

The recombination history of *L. aethiopica* was investigated by calculating the LD decay with PopLDdecay v3.41 [26] and the inbreeding coefficient *F*_IS_ after taking into account Wahlund effects. To this end, we considered five isolates (117-82, 1561-80, 169-83, 32-83 and 68-83) belonging to the same genetic population as inferred by ADMIXTURE and sampled over a period of four years. *F*_IS_ was calculated as 1-Ho/He, with Ho the observed heterozygosity and He the expected heterozygosity.

## RESULTS

### Genomic confirmation of interspecific hybridization including *L. aethiopica* as parent

Sequences were mapped against the *L. aethiopica* L147 reference genome, resulting in a median coverage of 27x to 70x for the publicly available sequence data, and 44x to 119x for the *L. aethiopica* isolates (Supp. Table 2). At least 80% of the positions in the reference genome were covered with at least 20 reads, and 72% of the paired reads aligned in proper pairs (Supp. Table 2). These results show that the coverage of the *L. aethiopica* reference genome was sufficiently large for variant calling, despite the diversity of species included in this study. Genotyping was performed with GATK HaplotypeCaller across all 36 *Leishmania* isolates, revealing a total of 988,363 high-quality bi-allelic SNPs.

A phylogenetic network revealed a total of four groups of isolates that corresponded to the four species included in this study (Figure 1A): *L. aethiopica* (20 isolates), *L. donovani* species complex including *L. donovani* (11 isolates) and *L. infantum* (1 isolate), *L. major* (1 isolate) and *L. tropica* (1 isolate). Two isolates did not cluster with any of these species: LEM3469 and its derived clones are positioned in between *L. aethiopica* and the *L. donovani* complex, and L86 is positioned in between *L. aethiopica* and *L. tropica* (Figure 1A). The network shows strong reticulation and a net-like pattern along the branches of these two isolates, suggesting that LEM3469 and L86 are interspecific hybrid parasites or the result of a mixed infection.

**Figure 1.**
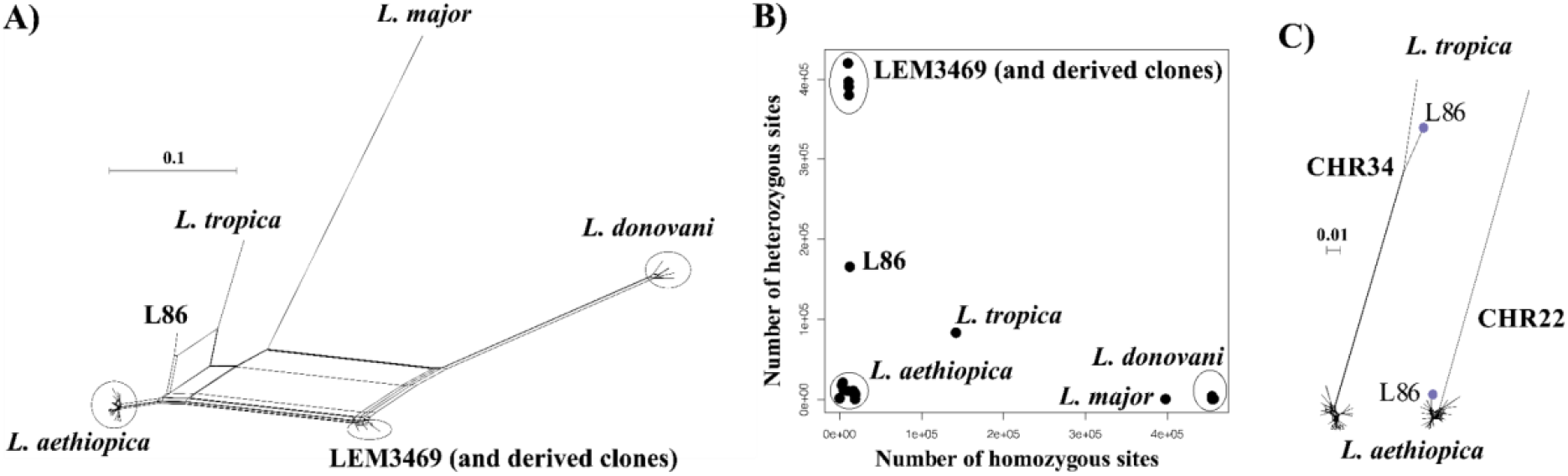
**A)** Phylogenetic network based on SNPs called across 36 genomes of *L. aethiopica*, the *L. donovani* species complex, *L. major* and *L. tropica*. **B)** Number of heterozygous sites versus number of homozygous sites for each of the 36 *Leishmania* genomes. **C)** Phylogenetic network based on SNPs called in the last 700kb of chromosome 34 and the first 300kb in chromosome 22. Blue dot indicates the position of the interspecific *L. aethiopica* - *L. tropica* hybrid L86.

For each genome, we counted the number of homozygous SNPs (where both haplotypes are different from the L147 reference) and heterozygous SNPs (where one haplotype is similar to the L147 reference and the other different). This showed that *L. donovani* (±414,160 SNPs), *L. major* (394,607 SNPs) and *L. tropica* (140,774 SNPs) genomes contain a relatively large amount of homozygous SNPs, and are thus genetically distant from *L. aethiopica* L147 (Figure 1B). In contrast, all *L. aethiopica* genomes and isolates L86 and LEM3469 (and derived clones) contained respectively 3,973 and ±12,230 homozygous SNPs (Figure 1B). The genomes of uncertain ancestry (L86 and LEM3469) displayed a much larger amount of heterozygous SNPs (164,566 in L87 and 377,135-416,672 in LEM3469) compared to the *L. aethiopica* genomes (422-20,783 SNPs) (Figure 1B). Such high levels of heterozygosity in L87 and LEM3469 further indicate the presence of divergent homologous chromosomes, either as the result of hybridization or because of a mixed infection.

When investigating the distribution of SNPs across the 36 chromosomes in LEM3469 and L86, we found that the majority of chromosomes consists almost entirely of heterozygous SNPs (Supp. Table 3). Exception was one genomic region in L86 (the last 700kb of chromosome 34) that was entirely homozygous for the alternate allele (both haplotypes are thus different from L147). In addition, the first 300kb of chromosome 22 in L86 and chromosomes 9, 10 and 15 in LEM3469 (Supp. Table 3) were virtually devoid of SNPs (both haplotypes are thus similar to L147). This observation of a largely heterozygous genome that is interrupted by homozygous stretches suggests that isolates L86 and LEM3469 are hybrid parasites, rather than the result of a mixed infection.

Similarly to LEM3469, all five derived clones of LEM3469 were SNP-poor in chromosomes 9, 10 and 15 (Supp. Figure 1, Supp. Table 3). All five clones were also SNP-poor in chromosomes 11 and 24, LEM3469 clones 1 and 9 were SNP-poor in chromosome 20,

LEM3469 clone 7 was SNP poor in chromosome 1 and LEM3469 clone 8 was SNP poor in chromosome 33 (Supp. Figure 1, Supp. Table 3). These results imply a loss of heterozygosity through the process of cloning and culturing of LEM3469.

Phylogenies based on SNPs in genomic regions that were either largely homozygous or SNP-poor in L86 revealed that this isolate clustered with either *L. tropica* or *L. aethiopica*, respectively (Figure 1C). This clearly points to *L. aethiopica* and *L. tropica* as the two parental species for L86. A phylogeny based on SNPs in chromosome 15 showed that LEM3469 and derived clones clustered within the *L. aethiopica* group (result not shown), confirming that *L. aethiopica* is one of the parental species of this hybrid isolate. No genomic regions were identified within the LEM3469 isolate or its clones that could reveal the other parental species.

In order to identify and confirm the other parental species of the hybrid parasites, we identified fixed SNPs specific to *L. tropica* (52,958 SNPs), *L. donovani / L. infantum* (294,890 SNPS) and *L. major* (253,458 SNPs). This revealed that 88% of the fixed SNPs specific to *L. tropica* were heterozygous in L86, compared to 0.3% of the *L. donovani* or *L. major*-specific SNPs. Similarly, we found that 91% of the fixed SNPs specific to *L. donovani* were heterozygous in LEM3469, compared to 0.01% of the *L. major* and *L. tropica-specific* SNPs.

Altogether, our results demonstrate that L86 and LEM3469 are the result of hybridization, rather than the result of a mixed infection, between *L. aethiopica* and *L. tropica* (in case of L86) or between *L. aethiopica* and *L. donovani* (in case of LEM3469).

### Population genomic structure and diversity of *L. aethiopica*

Variant calling was repeated on a dataset including solely the 20 *L. aethiopica* isolates, i.e. excluding the other Old World *Leishmania* species and the interspecific hybrids LEM3469 and L86. This resulted in a total of 94,581 high-quality INDELs and 284,776 high-quality SNPs called across 20 *L. aethiopica* genomes. Despite the relatively high genome-wide density of SNPs (median 89 SNPs and average 91 SNPs per 10 kb window), we found a relatively low number of heterozygous sites (median 6 SNPs and average 7 SNPs per 10 kb window). The allele frequency spectrum was dominated by low-frequency variants, with 58% of SNPs (170,163 SNPS) occurring at <0.1% of the population.

Our panel of *L. aethiopica* isolates displayed substantial differences in terms of the number of SNPs (5,607 - 93,762) and the number of homozygous (243 - 66,068) and heterozygous (2,067 - 69,857) SNP counts (Table 1). Most remarkable were i) isolate 1123-81 showing a low number of SNPs (5.607) compared to the other genomes (mean = 82,251; median = 87,132) and ii) isolates L100 and L100cl1 that were almost devoid of heterozygous SNPs (2,071) compared to the other genomes (mean = 32,251; median = 29,985). In addition, the number of fixed SNP differences between isolates ranged between 1 and 76,477 (average = 32,998 SNPs, median = 28,688 SNPs, sd = 20,955 SNPs) (Supp. Table 4). Exceptions were two groups of isolates that each contained two near-identical isolates (Supp. Table 4): i) strain 678-82 and LEM2358cl3 (a derived clone of the LEM2358 strain) were sampled eleven years from each other and were probably the result of clonal propagation in nature and ii) L100 and L100cl1, where L100 is a strain derived from a clinical sample and L100cl1 is a derived clone of the strain L100. Altogether, these results suggest that the *L. aethiopica* species is highly diverse and consists of both genetically similar and divergent parasites.

**Table 1.**
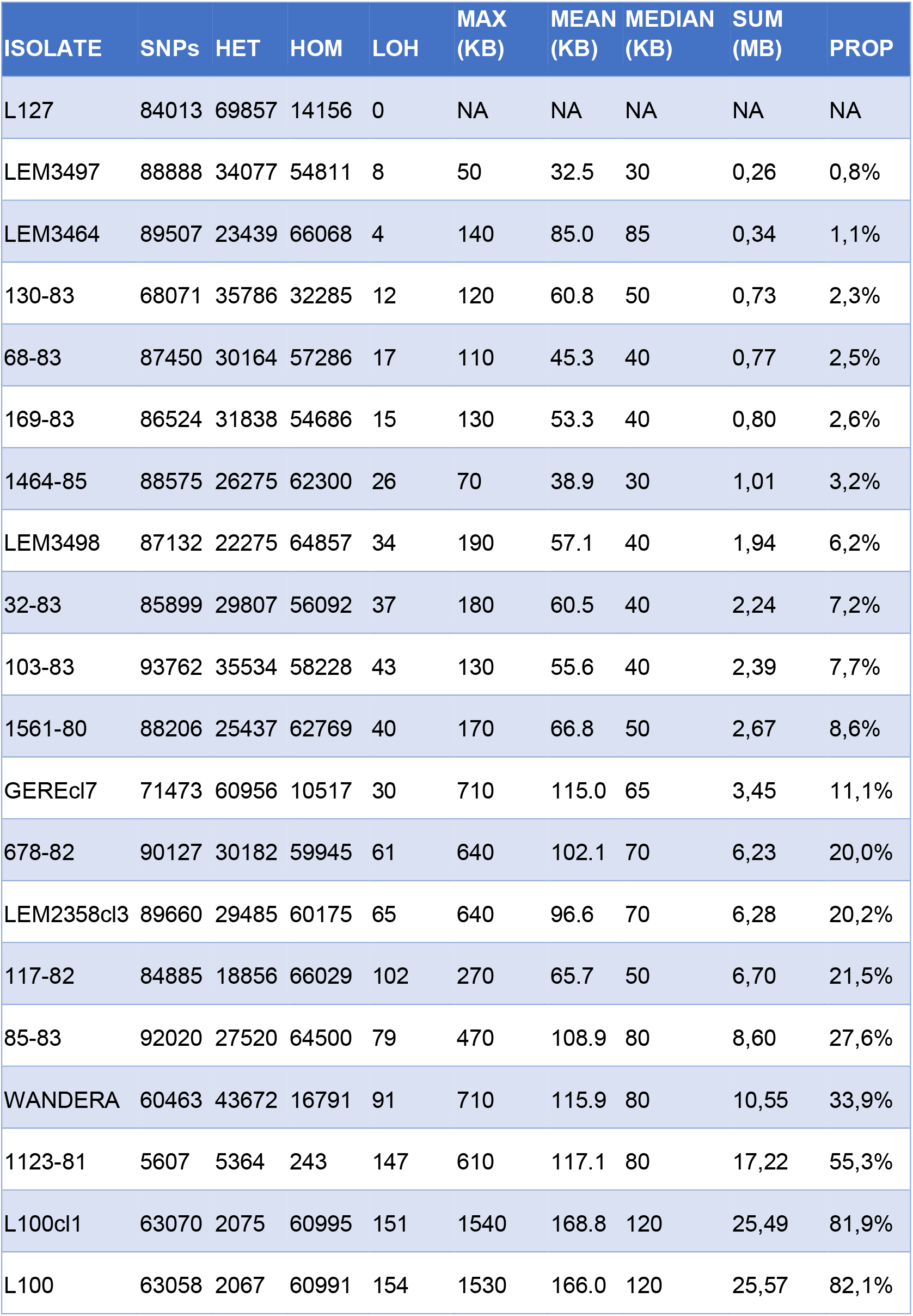
Population genomic statistics for each of the 20 *L. aethiopica* isolates: number of SNPs (SNPs), number of homozygous (HOM) and heterozygous (HET) SNPs, number of Loss-Of-Heterozygosity regions (LOH), maximum (MAX), mean (MEAN) and median (MEDIAN) length of LOH regions, total (SUM) and proportional (PROP) length of chromosomal genome covered by LOH regions.

The large number of fixed SNP differences prompted us to investigate the genome-wide distribution of Loss Of Heterozygosity (LOH) regions. This effort revealed major differences in the number and length of LOH regions between *L. aethiopica* isolates (Table 1) (Figure 2). The median length of LOH regions ranged between 30 kb in LEM3497 and 1464-85 to 120 kb in L100 and L100cl1 (Table 1). In particular isolates 1123-81, L100 and L100cl1 showed a high number of LOH regions (147 for 1123-81, 151 for L100 and 154 for L100cl1) covering a substantial proportion of their chromosomal genome (55.3% for 1123-81, 81.9% for L100 and 82.1% for L100cl1) (Table 1) (Figure 2). The largest LOH regions were at least 1.5 Mb long (Table 1) and were found in chromosome 36 of isolates L100 and L100cl1.

**Figure 2.**
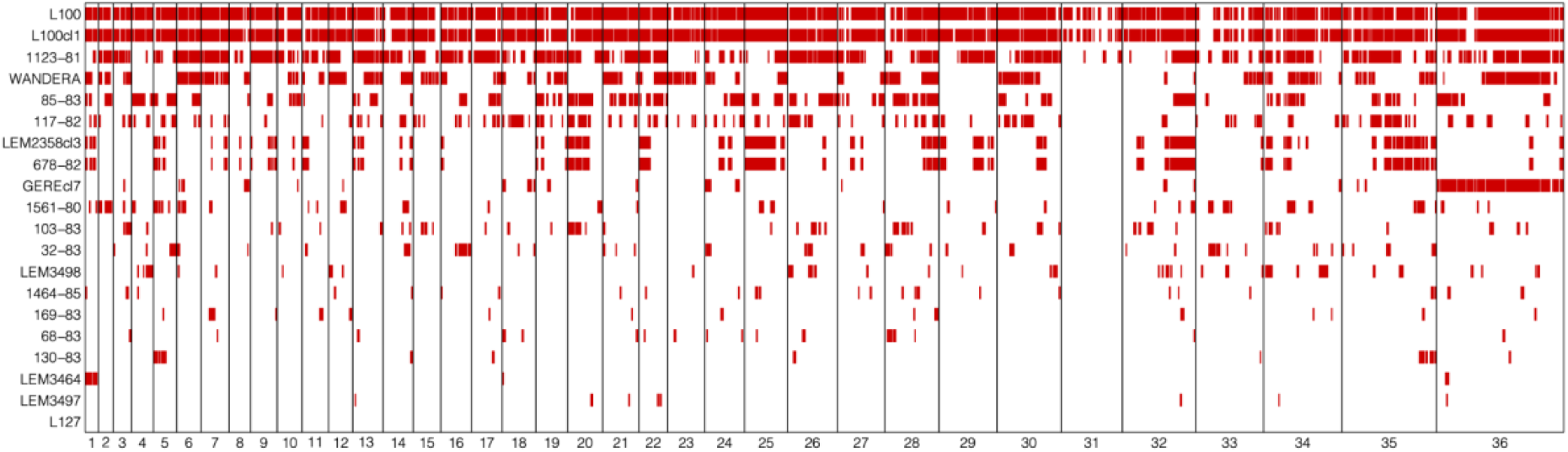
Loss-Of-Heterozygosity regions (red bars) in each of the 20 *L. aethiopica* isolates across the 36 chromosomes.

The population genomic diversity and structure of *L. aethiopica* was investigated after removing one individual from the near-identical isolate-pairs (678-82 and L100cl1) and excluding sites with high LD. ADMIXTURE analyses suggested the possible presence of two (K=2) to three (K=3) ancestral populations within our panel of *L. aethiopica* isolates, although the 5-fold cross validation was approximately similar for *K*=1 (Supp. Figure 2). The ancestry estimation for the different values of *K* were consistent over the different SNP-pruning thresholds (Supp. Figure 2). Assuming *K*=3 populations, all but two isolates (GEREcl7 and WANDERA) were assigned to one of the three ancestry components with >99% probability (Figure 3A). Principal Component Analysis (PCA) (Supp. Figure 3), a phylogenetic network based on genome-wide SNPs (Figure 3B) and phylogenetic networks based on SNPs in LOH-poor chromosomes (Supp. Figure 4) showed a clear separation among groups of individuals corresponding to the clusters inferred by ADMIXTURE. These results show congruence in *L. aethiopica* population structure among various inference approaches and suggest that the presence of LOH regions has little impact on the inference process (as shown in Supp. Figure 4). Mean pairwise *F*_ST_ values among these three populations revealed strong population differentiation, ranging between 0.175 and 0.235 (Supp. Table 5).

**Figure 3.**
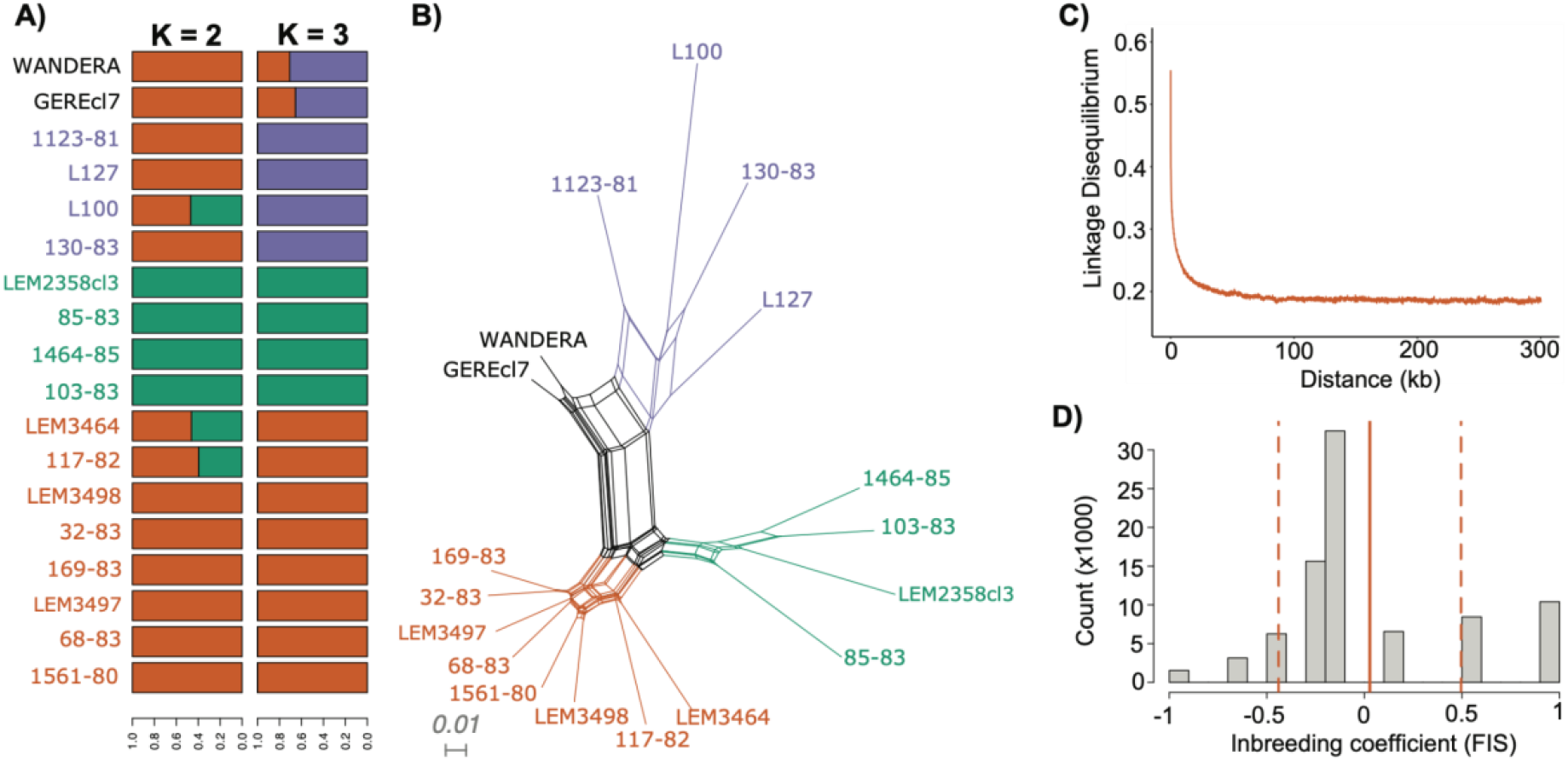
Population genomic diversity and structure of *L. aethiopica*. **A)** Barplots depicting ancestral components as inferred by ADMIXTURE for K = 2 and K = 3 populations, based on 85,725 SNPs. **B)** Phylogenetic network based on uncorrected p-distances among 18 *L. aethiopica* genomes genotyped at 277,156 bi-allelic SNPs. Coloured branches and tip labels correspond to the inferred populations by ADMIXTURE at K=3. **C)** Linkage decay plot for 1561-80, 68-83, 169-83, 32-83 and 117-82 controlling for spatio-temporal Wahlund effects (see methods). **D)** *F* is distribution after correction for spatio-temporal Wahlund effects for 1561-80, 68-83, 169-83, 32-83 and 117-82. The solid line represents the mean *F* is values (0.027) while the dashed lines represent the standard deviation (± 0.47).

The phylogenetic network also revealed a reticulated pattern and long terminal branches, indicative of recombination (Figure 3B). Estimates of relative recombination rates in *L. aethiopica* were calculated by *F*_IS_ per site and LD decay controlling for spatio-temporal Wahlund effects (see methods) (Figure 3C, D). The per-site *F*_IS_ was unimodally distributed and close to zero (mean *F*_IS_ = 0.027 ± 0.47). In addition, LD levels were relatively low with r^2^ descending to 0.2 at 21.9kb. These results suggest relatively frequent genetic exchange in *L. aethiopica*.

### Chromosomal and local copy number variations in *L. aethiopica*

The ploidy of all isolates was investigated using the genome-wide distribution of alternate allele read depth frequencies at heterozygous sites (ARDF), which should be centered around 0.5 in diploid organisms. The genome-wide distribution of ARDF was unimodal and centered around 0.5 for all *L. aethiopica* isolates and the *L. aethiopica - L. tropica* hybrid L86, suggesting that the baseline ploidy of these parasites is diploid. However, the distribution of ARDF was bimodal with modes 0.33 and 0.67 for the *L. aethiopica* - *L. donovani* hybrid LEM3469 and its derived clones, suggesting that the baseline ploidy of this hybrid is triploid.

Variation in chromosomal copy numbers was further investigated using normalized chromosomal read depths (RD). The RD estimates revealed that chromosome 31 was at least tetrasomic in all isolates (Figure 4). Little variation was detected for six *L. aethiopica* isolates that were largely diploid (box 1 in Figure 4). The rest of the *L. aethiopica* isolates were trisomic at 1 to 6 chromosomes, including a group of 8 isolates that was trisomic for chromosome 1 (box 2 in Figure 4). The largest variability was found in *L. aethiopica* isolate LEM3498, the *L. aethiopica* - *L. donovani* hybrid LEM3469 and its derived clones and the *L. aethiopica* - *L. tropica* hybrid L87 (box 3 in Figure 4). Altogether, our results demonstrate that *L. aethiopica* genomes are aneuploid, as shown for other *Leishmania* species. Moreover, note that we also observed non-integer values of somy for some chromosomes (Figure 4), suggesting the presence of mosaic aneuploidy [26].

**Figure 4.**
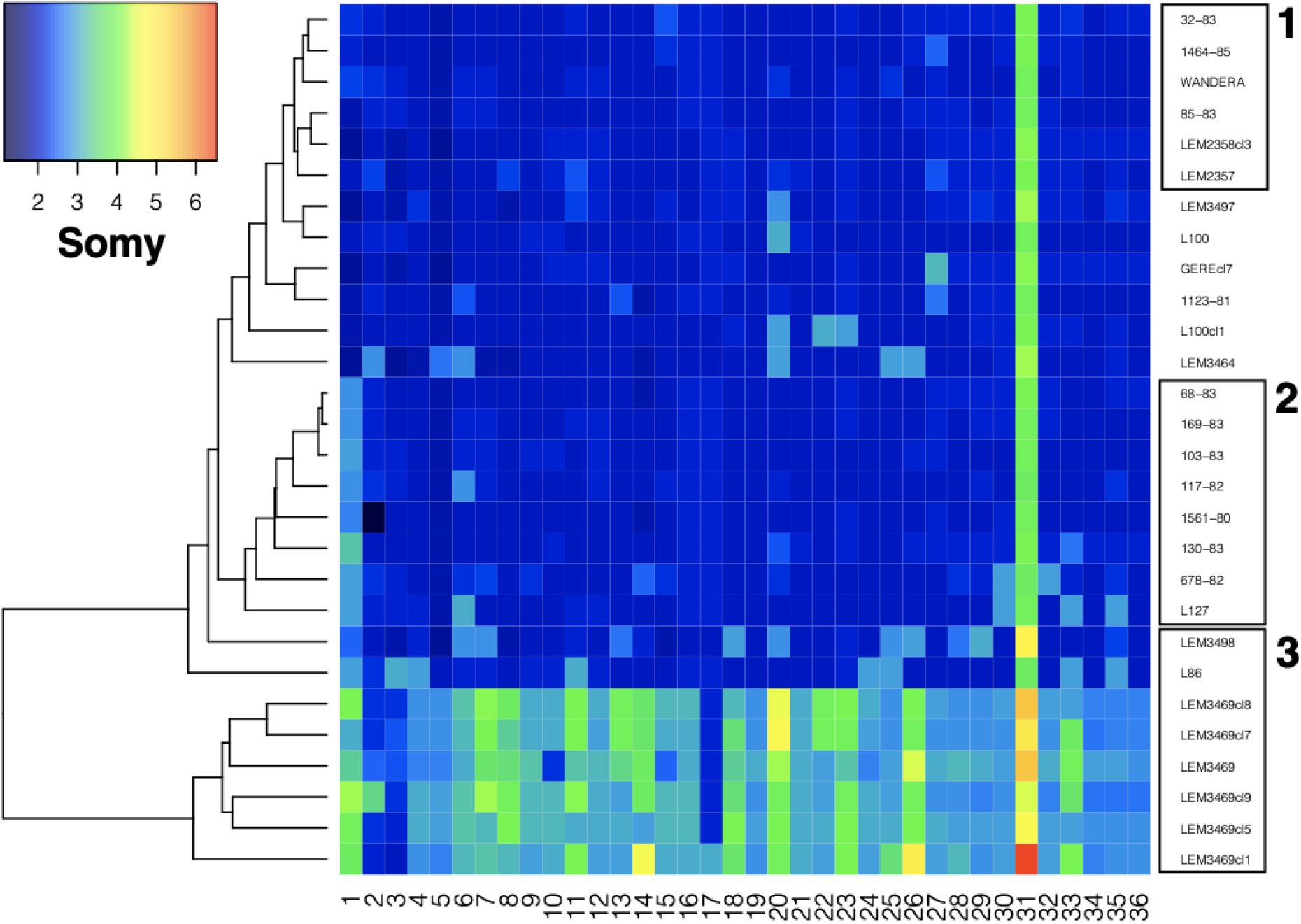
Somy variation across 36 chromosomes for the 28 *Leishmania* genomes sequenced in this study. Box 1 highlights isolates that are nearly diploid, box 2 highlights isolates with a trisomic chromosome 1 and box 3 highlights isolates showing high somy variability (see text).

Local copy number variations were investigated using normalized haploid copy numbers (HCN) as estimated in non-overlapping 2kb windows. We identified a total of 379 genomic regions with decreased or amplified HCN across our panel of *L. aethiopica* isolates. The majority of these windows were only 2kb long (N = 266; 70%) and the mean (1.1 HCN) and median (0.79 HCN) differences in HCN across our panel were minor, suggesting that our sample of *L. aethiopica* displays relatively little local copy number variation. Apart from this observation, most notable was a 46 kb telomeric genomic region on chromosome 29 in isolate 1561-80 showing a 10-fold increase in HCN compared to the other *L. aethiopica* isolates. This region covers a total of 18 protein coding genes, including a ribonuclease inhibitor-like protein (LAEL147_000545200), an actin-like protein (LAEL147_000544400) and a putative inosine-adenosine-guanosine-nucleoside hydrolase (LAEL147_000545100) (Supp. Table 6). Several other isolates (L127, GEREcl7, 169.83 and 32.83) displayed intrachromosomal amplifications across genomic regions of 10 kb to 20 kb long, but these involved only 0.3 to 1.2 increase in HCN (Supp. Table 6). Finally, several small (<= 4kb) intrachromosomal amplifications were identified across the genomes of isolates L100 and L100cl1 (Supp. Table 6).

Within the *L. aethiopica* - *L. donovani* hybrid LEM3469 and its derived clones, we detected a 6kb genomic region in chromosome 29 showing a 2-fold increase in HCN and covering a Leucine Rich Repeat gene (LAEL147_000538100) (Supp. Table 7). Within the *L. aethiopica* - *L. tropica* hybrid L87, we found a 130 kb genomic region in chromosome 35 with a 1.6 fold increase in HCN and covering a total of 46 protein coding genes (Supp. Table 8). Within both hybrids, we detected a 14 kb - 16 kb genomic region in chromosome 30 showing a ~1-fold increase in HCN and covering several protein coding genes such as a the putative ferric reductase gene (LAEL147_000563600) (Supp. Tables 7-8).

## DISCUSSION

We present the first comprehensive genome diversity study of *L. aethiopica* by analyzing high-resolution WGS data. This revealed insights into the genetic consequences of recombination in *Leishmania* at both the species- and population-level.

At the species level, we provide genomic evidence of hybridization involving *L. aethiopica* as one of the parental species and *L. tropica* (in case of the L86 hybrid strain) or *L. donovani* (in case of the LEM3469 hybrid strain) as the other parental species. Strain LEM3469 has already been described as a potential *L. aethiopica /L. donovani* hybrid based on microsatellite data and single-gene sequences [13]. Strain L86 has been flagged as a potential hybrid, although the isolate remained classified as *L. aethiopica* due to lack of molecular evidence [27]. Here, our genome analyses provide unequivocal evidence that these two *Leishmania* strains are interspecific hybrids. Our finding of genome-wide heterozygosity that is only occasionally interrupted by patches of homozygosity suggest that these two hybrids are equivalent to F1 progeny that propagated mitotically since the initial hybridization event. Similar genomic descriptions of naturally circulating F1-like hybrids were advanced previously for *L. braziliensis* x *L. peruviana* in Peru [8] and for *L. braziliensis* x *L. guyanensis* in Costa Rica [28]. Our results thus support a growing body of genomic evidence for extensive genetic exchange in protozoan parasites in the wild [7–9,28–33].

The *L. tropica* / *L aethiopica* (L86) hybrid strain was near diploid, being disomic at 27/36 chromosomes, suggesting balanced segregation of the chromosomes during hybridization. Trisomic chromosomes in L86 contained either one or two copies of the *L. tropica* parent, suggesting that these trisomies arose after the hybridization event through multiplication of one of the parental chromosomes, possibly due to culturing *in vitro* [34]. These observations seem unlikely to have arisen by a random parasexual process, and strongly suggest that the L86 hybrid is the result of meiotic recombination. The *L. donovani* / *L aethiopica* (LEM3469) hybrid strain was near triploid, being trisomic at 21/36 chromosomes. Triploid hybrids have been routinely recovered from experimental matings in *Leishmania* [9] and from clinical samples of cutaneous leishmaniasis patients [28], and are observed across a variety of organisms capable of sexual reproduction [35–37]. Results from experimental crosses in *Leishmania* suggested that interspecific hybrids with close to 3n DNA content were likely due to an asymmetric meiosis between a parental 2n cell that failed to undergo meiosis and 1n gamete from the other parent [9], similar to what has been suggested for *T. brucei* [38]. One observation that requires further investigation is that triploid hybrids may occur more frequently when the two parental species are genetically divergent (in the case of *L. donovani* / *L aethiopica*, *L. guyanensis* / *L. braziliensis* [28] and *L. infantum* / *L. major* [9] hybrids), while diploid hybrids seem to occur when the parental species are more closely related (in the case of *L. tropica* / *L. aethiopica* and *L. braziliensis* / *L. peruviana* [8] hybrids).

At the population level, our analyses of sequence variation confirmed previous observations that the *L. aethiopica* species is genetically diverse, despite its restricted geographic distribution [10–12]. Indeed, we detected on average 91 SNPs per 10kb in *L. aethiopica*, which is comparable to the genetically diverse *L. braziliensis* (106 SNPs per 10kb) but twice the number observed in the demographically bottlenecked *L. peruviana* (41 SNPs per 10kb) [8]. By comparison, the human genome contains ~8 SNPs per 10kb [39]. Also, the total number of SNPs (284,776) across our set of 20 *L. aethiopica* genomes from Ethiopia is of the same magnitude as the number of SNPs (395,624) observed in a set of 151 globally sampled genomes of the *L. donovani* species complex [40]. In contrast, we detected relatively little variation in local copy numbers: the majority of isolates contained only up to 20 genomic regions with increased/decreased read depths, and most of these genomic regions were only 2 kb long. Hence, adaptation in our sample of *L. aethiopica* isolates may depend on sequence variation rather than gene dosage. Future studies involving direct sequencing of biopsy samples [41] should allow deciphering the relative contribution of different genomic variants in clinical outcome or treatment failure.

Population genomic analyses revealed that the *L. aethiopica* species consists of both asexually evolving strains (as indicated by the existence of near-identical isolates) and groups of parasites that show signatures of recombination (as indicated by linkage decay). In addition, parasite populations were genetically strongly differentiated, suggesting that *L. aethiopica* consists of divergent ancestral populations, although lack of geographic/ecological data precluded us from studying its evolutionary history in greater detail. A remarkable observation was the extensive loss of heterozygosity (LOH) in some *L. aethiopica* strains. Such LOH likely arose from gene conversion/mitotic recombination and indicate that *L. aethiopica* strains may evolve asexually over relatively long time periods. This is exemplified by two near-identical genomes (678-82 and LEM2358cl3) that were sampled nine years from each other. Interestingly, we observed four additional LOH regions in LEM2358cl3 (sampled in 1991) impacting an additional 50 kb of the chromosomal genome compared to 678-82 (sampled in 1982), suggesting that nine years of clonal evolution has resulted in LOH across 50kb of the genome. Knowing that 6.28 Mb of the LEM2358cl3 genome is impacted by LOH, we extrapolate that the original strain may have propagated mitotically for roughly 1,200 years.

Extensive LOH regions have been described previously for obligate asexual eukaryotes, such as the water flea *Daphnia pulex* [42] and the protozoan parasite *Trypanosoma brucei gambiense* [25,43]. In *T. b. gambiense*, most LOH regions are ancestral and thus present in all strains, or at least within sets of strains that share a common ancestry [25]. In contrast, we observed large differences in the number and length of LOH regions between *L. aethiopica* strains, irrespective of their ancestral relationships. For instance, LOH regions were absent in the L127 strain while a total of 154 LOH regions were found in the L100 strain, impacting at least 82% of its chromosomal genome. The L100 strain (also known as LEM144) was isolated in 1972 and has - to the best of our knowledge - only been cultured *in vitro* for about 30 passages before sequencing. Hence, the extensive loss of heterozygosity can not be explained by maintenance *in vitro*, and is probably due to long-term mitotic recombination in the wild, as described for the asexually evolving *T. b. gambiense* [25]. Alternatively, the genome-wide loss-of-heterozygosity in L100 may be explained by genome-wide DNA haploidization, possibly due to the production of haploid gametes as seen in trypanosomes [44], followed by whole-genome diploidization (as our analyses demonstrated that L100 is diploid). However, there is currently no evidence in literature for the occurrence of whole-genome haploidization/diploidization in *Leishmania* parasites.

In conclusion, our study has shown that *L. aethiopica* is genetically diverse and divided into divergent populations, suggesting that strains may be ecologically/geographically confined. While linkage decay suggests the occurrence of genetic exchange, the discovery of extensive loss of heterozygosity also suggests that some *L. aethiopica* strains may evolve asexually for relatively long periods of time. Hence, *L. aethiopica* presents an ideal model to understand the impact of parasite population structure and hybridization on genome evolution in protozoan parasites. Our preliminary observations should thus be investigated in further detail using larger sets of isolates/samples for which detailed eco-epidemiological data are available, preferably using direct-sequencing approaches that would also allow understanding the genomic basis of clinical and treatment outcome.

## ACKNOWLEDGEMENTS

This work was supported by the Dioraphte foundation (Spatial-CL, project number: CFP-RD2020 20020401) and the Belgian Directorate General for Development Cooperation (fourth framework agreement, project 907004). FVDB and SH acknowledge support from the Research Foundation Flanders (Grants 1226120N and G092921N).

## DATA AND SCRIPT AVAILABILITY

Genomic sequence reads of all sequenced isolates will be made available on the European Nucleotide Archive (www.ebi.ac.uk/ena) upon publication of this manuscript. Unless specified in the text, scripts are available at https://github.com/amberhadermann/L.aethiopica.

---

**Supplementary Figure 1.**
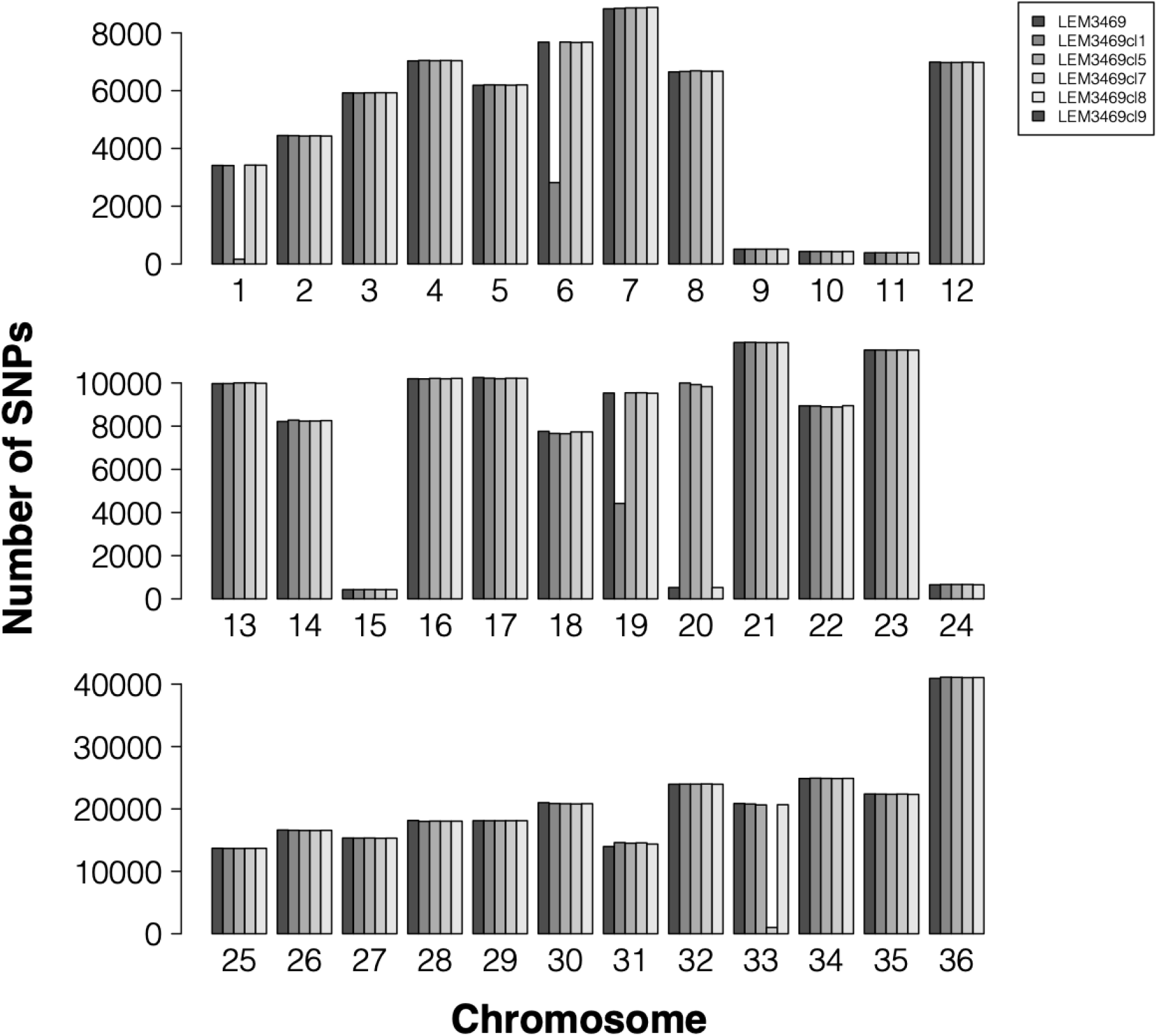
Number of SNPs for each of the 36 chromosomes in LEM3469 and its five derived clones.

**Supplementary Figure 2.**
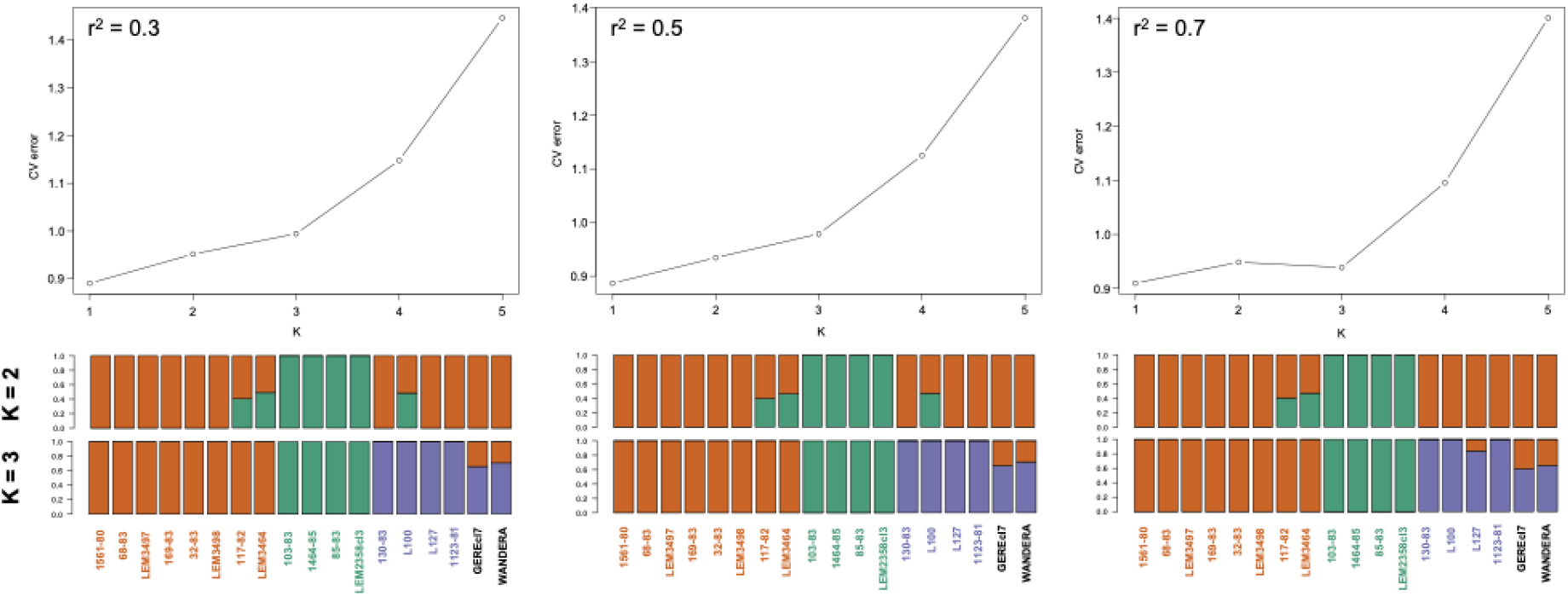
Model-based ancestry estimation of *L. aethiopica*, as inferred by ADMIXTURE, for different SNP-pruning thresholds (see methods). Upper panels depict the 5-fold cross validation plots for K = 1-5. Lower panels represent barplots of the ancestral components inferred by ADMIXTURE for K = 2 and K = 3. (left) SNP pruning at r^2^=0.3 retaining 47,244 SNPs. (middle) SNP pruning at r^2^=0.5 retaining 85,725 SNPs. (right) SNP pruning at r^2^=0.7 retaining 112,241 SNPs.

**Supplementary Figure 3.**
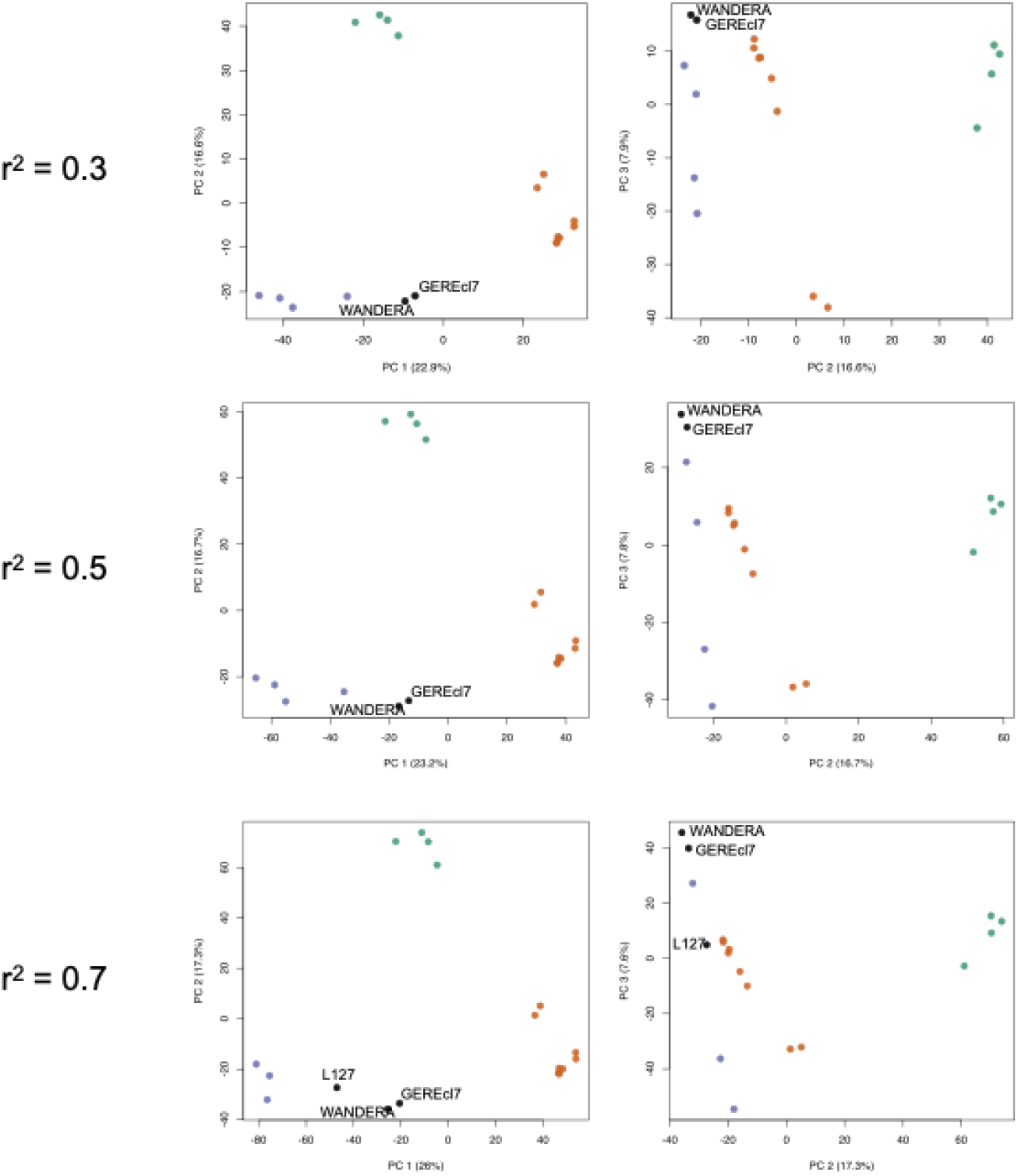
Principal component analysis for the different SNP-pruning thresholds (see methods). (upper) SNP pruning at r^2^=0.3 retaining 47,244 SNPs. (middle) SNP pruning at r^2^=0.5 retaining 85,725 SNPs. (lower) SNP pruning at r^2^=0.7 retaining 112,241 SNPs. Colors represent the population assignment as inferred by ADMIXTURE (Supplementary Figure 3).

**Supplementary Figure 4.**
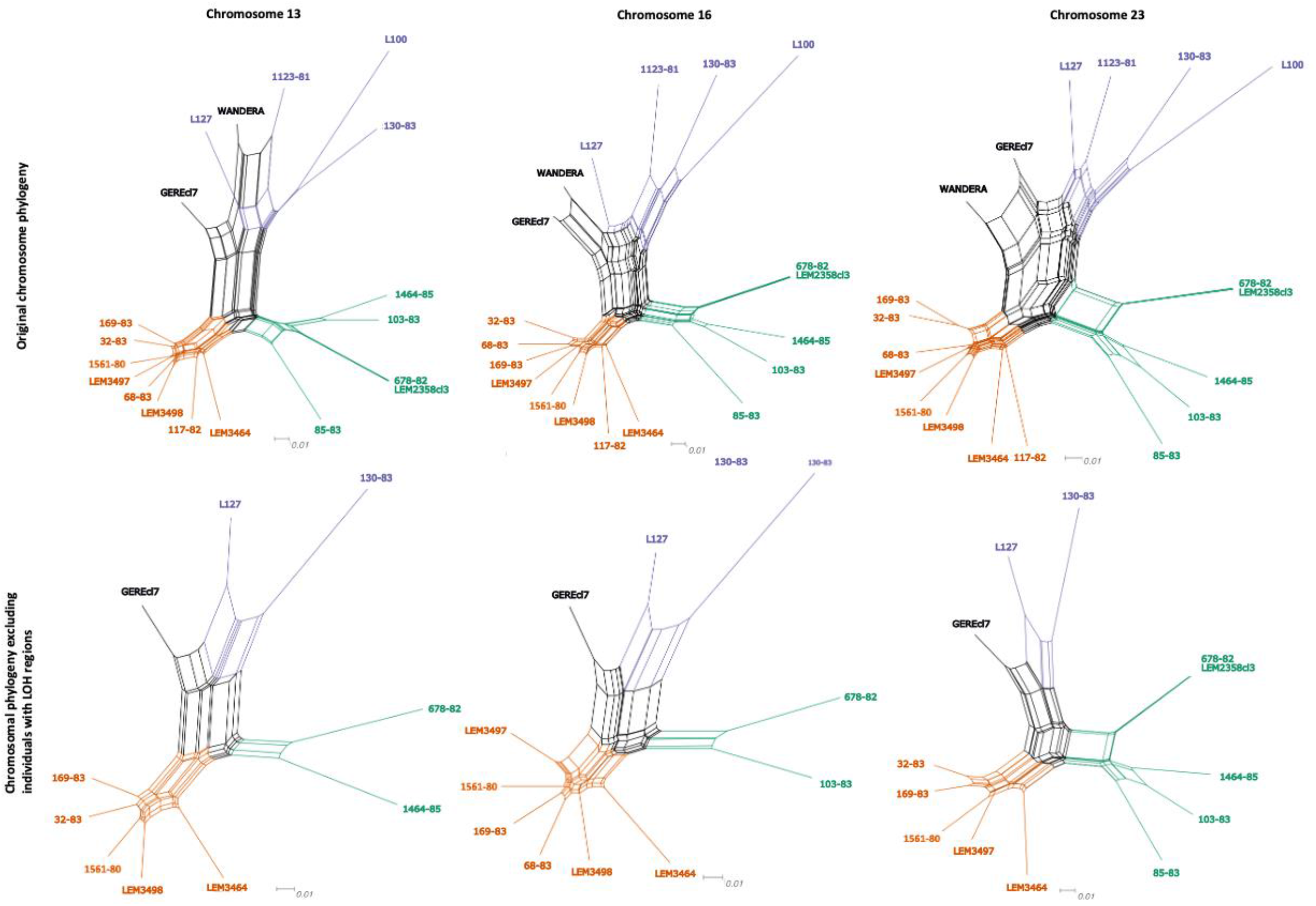
Phylogenetic network based on uncorrected p-distances for chromosome 13 (left), 16 (middle) and 23 (right) with (lower) and without (upper) excluding individuals containing LOH regions. Coloured branches and tip labels correspond to the inferred populations by ADMIXTURE at K=3 (Figure 3A).

